# Multinucleation Associated DNA Damage causes quiescence despite compromised p53

**DOI:** 10.1101/2020.12.22.424035

**Authors:** Madeleine Hart, Sophie D Adams, Viji M Draviam

**Affiliations:** School of Biological and Chemical Sciences, Queen Mary University of London, London, UK

**Author notes:** Address correspondence to: Viji M Draviam, School of Biological and Chemical Sciences, Queen Mary University of London, London E1 4NS.

## Abstract

Nuclear atypia is one of the earliest hallmarks of cancer progression. How distinct forms of nuclear atypia differently impact cell fate is not understood at the molecular level. Here, we perform single-cell tracking studies to determine the immediate and long-term impact of multinucleation or misshapen nuclei and reveal a significant difference between multinucleation and micronucleation, a catastrophic nuclear atypia known to promote genomic rearrangements and tumour heterogeneity. Tracking the fate of newborn cells exhibiting various nuclear atypia shows that multinucleation, unlike other forms of nuclear atypia, blocks proliferation in p53-compromised cells. Because compromised p53 is seen in over 50% of cancers, we explored how multinucleation blocks proliferation and promotes quiescence. Multinucleation increases 53BP1-decorated nuclear bodies (DNA damage repair platforms), along with a heterogeneous reduction in transcription and protein accumulation across the multi-nucleated compartments. Importantly, Multinucleation Associated DNA Damage (MADD) associated 53BP1-bodies remain unresolved for days, despite an intact NHEJ machinery that repairs laser-induced DNA damage within minutes. This persistent MADD signalling blocks the onset of DNA replication and is associated with driving proliferative G1 cells into quiescence, revealing a novel replication stress independent cell cycle arrest caused by mitotic lesions. These findings call for segregating protective and prohibitive nuclear atypia to inform therapeutic approaches aimed at limiting tumour heterogeneity.

## INTRODUCTION

Nuclear atypia is associated with disease states, and is a strong discriminator of survival in many cancers^1,2^. Micronuclei harbour nuclear envelope defects^3^, leading to cell cycle asynchronicity between primary and micronucleus^4^ and extensive DNA damage following replication stress^4,5^. Furthermore, the erroneous repair of the micronuclear DNA^5,6^ promotes large scale translocations confined to micronuclear DNA^5^. Thus, micronuclei can propagate genomic instability and therefore, considered tumourigenic, particularly given their ability to proliferate in p53 null conditions^4,7^. While micronuclei studies provide evidence that nuclear atypia may be causal to a disease state, what impact other nuclear atypia have upon a disease state is largely unknown.

Multinucleation, another form of nuclear atypia with multiple nuclear compartments, has been linked to mitotic slippage^8^ the erroneous exit from a prolonged mitotic arrest. Whilst multinucleated cells are known to display DNA damage^8–10^, their cell cycle fate compared to micronucleated cells and the dynamics of DNA damage and downstream signalling have not been assessed.

Mitotic errors can lead to a variety of nuclear atypia, including micronucleation, multinucleation or misshapen nuclei, arising either as a result of mitotic slippage (slow exit) or spindle checkpoint failure (abrupt exit) (reviewed in ^11^). While structural aneuploidies can impair cell cycle progression,^12^ the cell cycle impact of various nuclear atypia has not been compared so far within a common mitotic lesion. A molecular understanding how nuclear atypia impacts cell viability and DNA repair status in single-cells of a population is important to understand disease outcome heterogeneity.

## RESULTS

### Unlike other nuclear atypia, multinucleation arrests cells at G1-S, independent of p53

To mimic mitotic drug treatment associated nuclear atypia, we exposed cells to GSK-923295, a CENP-E inhibitor (CENPEi), which disrupts the end-on conversion of chromosome-microtubule attachment^13,14^ leading to chromosome misalignment and a mitotic arrest^15^ causing mitotic slippage and a variety of nuclear atypia. To analyse cell cycle status, we immunostained for PCNA foci (S-phase marker) two days after the release from CENPEi treatment (Fig. 1A). Using immunostaining we compared 100s of RPE1 cells displaying a variety of nuclear atypia. As expected, multinucleation, micronucleation or misshapen nuclei were all observed predominantly in the CENPEi-treated but not DMSO-treated population (Fig. 1B and 1C). Only in cells displaying multinucleation, no PCNA foci was observed; cells displaying micronuclei or normal or misshapen nuclei presented a small proportion of PCNA foci positive cells (Fig. 1B and 1D), showing a multinucleation associated block in G1-S transition and replication initiation.

**Figure 1 -.**
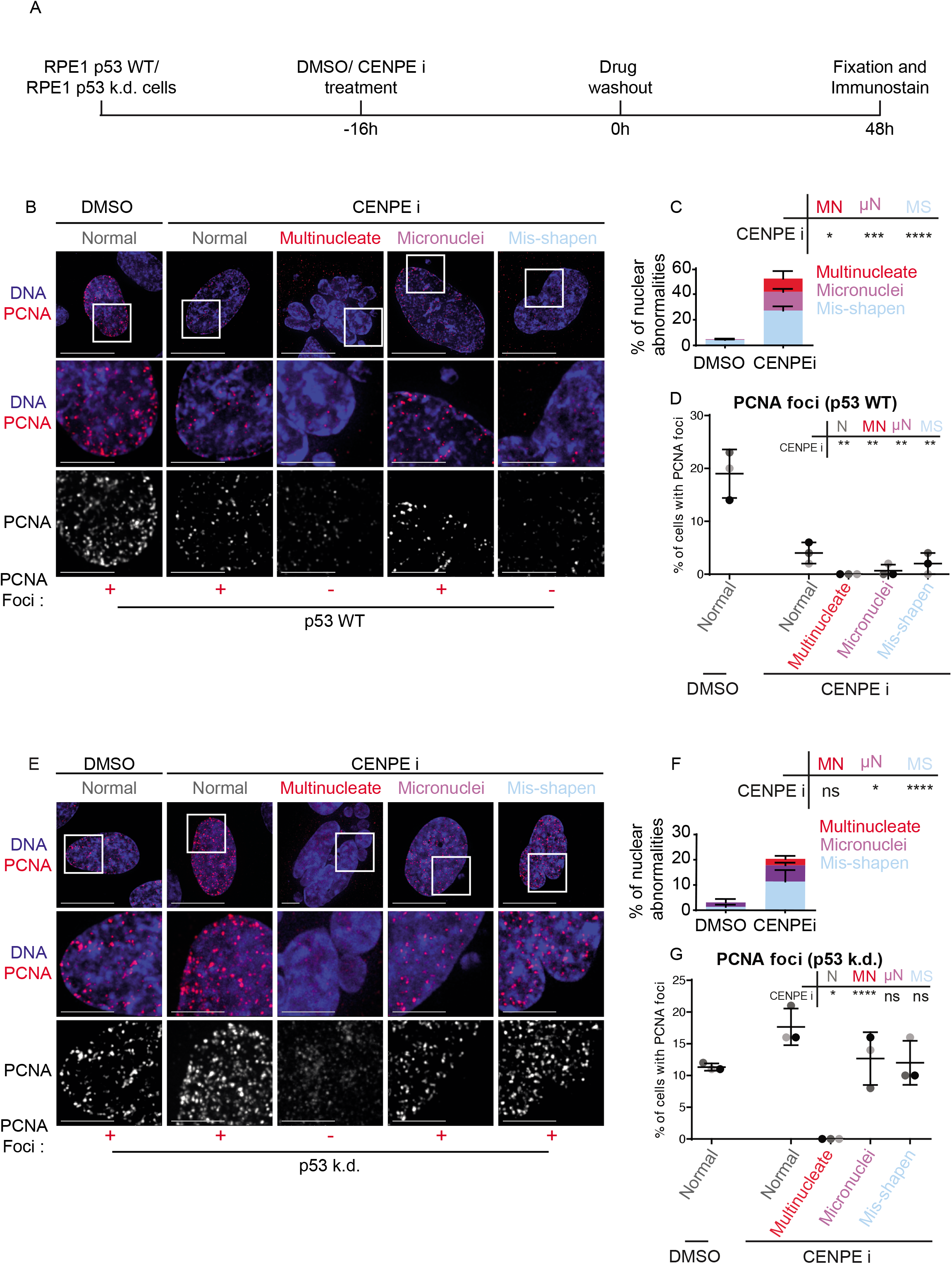
Unlike other nuclear atypia, multinucleation arrests cells at G1-S, independent of p53. **A** - Experimental regime; RPE1 p53 wild type (WT) or RPE1 p53 knockdown (k.d.) were treated with DMSO or CENPE inhibitor for 16 hours, then washed out. 48 hours later cells were assessed for PCNA and nuclear status. **B & E** - Images of RPE1 p53 WT (B) or p53 k.d (E) cells treated as in A. Scale 15μm, insets 5μm. +/- refers to PCNA foci positive or negative nuclei displayed. Note in **B a** micronuclei bearing cell displaying PCNA-foci has been chosen, although the majority lack PCNA-foci. **C & F**-Quantification of nuclear morphology changes after treatment of RPE1 p53 WT (C) or p53 k.d (F) cells with DMSO or CENPE inhibitor, as in A. Nuclei were classified as either Mis-shapen (MS), Micronucleated (μN) or Multinucleated (MN). N=600 cells, 3 independent experimental repeats shown as shades of grey. Statistical analysis was using a two-way ANOVA with multiple comparisons and a confidence interval of 95%. **D & G**-Quantification of the percentage of PCNA-foci positive cells, within each nuclear morphology bin, following DMSO or CENPE inhibitor treatment, of RPE1 p53 WT (C) or p53 k.d (F) cells. N>150, 3 independent experimental repeats shown as shades of grey. Statistical analysis was using multiple unpaired t-tests, comparing each morphology after CENPE inhibition to normal nuclei after DMSO treatment.

The complete absence of PCNA foci in multinucleated cells is somewhat at odds with reports of a positive association between multinucleation and tumorigenic potential and resistance to therapy^16,17^. So, we investigated whether the G1-S block in multinucleated cells may be lost following compromised p53 function, a tumour suppressor frequently lost in cancers (reviewed by ^18^). A p53 dependent G1-S block has been reported in micronucleated cells^7^, tetraploid cells^19^ and chromosomal missegregation or aneuploidy^7,20,21^, but not tested in multinucleated cells. We exposed RPE1 p53 knockdown cell line, with demonstrably no p53 expression^7^ (Supplementary Figure 1A, 1B) to CENP-E inhibitor and monitored PCNA status 48 hours after inhibitor wash-off (Fig. 1A). In p53 knockdown cells, where all three forms of nuclear atypia could be observed (Fig. 1F), nuclear PCNA foci, distinctly brighter than non-specific cytoplasmic speckles, were absent in multinucleated cells alone (Fig. 1E and 1G). In contrast, cells presenting other nuclear atypia, including micronucleated or misshapen nuclei showed an increase in the proportion of PCNA foci in p53 knockdown cells compared to WT RPE1 cells (Fig. 1E and 1G). The lack of PCNA foci selectively in multinucleated cells shows that unlike other forms of nuclear atypia, multinucleation blocks G1-S transition, independent of p53 status.

Two other mitotic lesions where multinucleation is not readily induced due to accelerated mitotic exit, inhibitions of Mps1 or Aurora-B activity, yielded similar results (Supplementary Fig. 2A, 2B). Assessing the cell cycle impact of micronuclei and misshapen nuclei showed that in p53 WT conditions both treatments exhibit a G1 arrest, but less robust than multinucleate cells (Supplementary Fig. 2C). Additionally, in p53 knockdown cells, the G1 arrest was alleviated in micronucleated and misshapen nuclei caused following MPS1 or Aurora-B inhibition (Supplementary Fig. 2D, 2F). In summary, unlike other nuclear atypia, multinucleation uniquely blocks G1-S despite a compromised p53 status.

### Multinucleate cells exhibit the highest incidence of DNA damage

We compared the extent of DNA damage in the various nuclear atypia, by immunostaining for gamma H2AX (double-stranded DNA damage marker), 1 and 48 hours after CENPEi washout. 48 hours after washout, 40% of micronuclei bearing cells displayed gH2AX foci exclusively within the micronuclei compartment, as reported^4,5,22^ and only 30% of their primary nuclei presented gH2AX foci (Supplementary Fig. 3A and 3B). In contrast, >90% of multinucleated cells displayed gH2AX foci in most of the compartments (Supplementary Fig. 3A and 3B). Importantly, the number of gH2AX foci is strikingly higher after multinucleation compared to other forms of nuclear atypia (Supplementary Fig. 3C), although the intensity of individual gH2AX foci per se was not different (Supplementary Fig. 3D). Similarly, within an hour of drug washout, gH2AX foci number was strikingly higher in multinucleated cells (Supplementary Fig. 4, see 4E). These observations show that multinucleated cells are prone to a substantially higher incidence of DNA damage compared to other forms of nuclear atypia.

Next, we analysed the incidence of RIF1, an NHEJ repair pathway member that accumulates at DNA breaks specifically in G1 phase^23–25^. Immunostaining studies showed the normal accumulation of RIF1 in the vast majority of gH2AX foci in multinucleated cells, except for the gH2AX foci in micronucleated compartments (Supplementary Figure 3F). We conclude that Multinucleation associated DNA damage (MADD) successfully accumulates RIF1, a factor that promotes NHEJ in G1.

### Multinucleate cells exhibit delayed DNA damage signalling

To identify the precise timing of MADD and the extent of MADD resolution, we used time-lapse microscopy to track the arrival and departure of 53BP1 foci, a DNA damage response factor downstream of gH2AX and upstream of RIF1. In RPE1 cells coexpressing 53BP1-GFP and Histone-2B-GFP (DNA marker), we compared multinucleated cells arising from a slow mitotic slippage or rapid mitotic exit following either CENP-Ei treatment alone or CENP-Ei and Aurora-Bi co-treatment, respectively (Supplementary Fig. 5A; Fig. 2A and 2B; Supplementary Videos 1A-C). Videos of CENPEi washout showed nearly 80% of cells exiting mitosis with a multinucleate phenotype and only 20% of cells exhibiting either a normal or misshapen nuclei (Supplementary Fig. 5B). 53BP1 foci could be observed in multinucleated cells, as well as those with normal or misshapen nuclei (Fig. 2C), consistent with gH2AX status (Supplementary Fig. 4), with multinucleated cells gaining foci in ~60% of cells during the course of imaging irrespective of slow mitotic slippage (CenpEi) or a rapid mitotic exit (CenpEi & AuroraBi). Analysis of live-cell videos revealed that the timing of 53BP1 foci arrival varies depending on the nuclear atypia: in multinucleated cells, 53BP1 foci arrival is delayed, ~2-11 hours after mitotic exit (Fig. 2D), whereas in others foci arrival was complete, within the first 4 hours after mitotic exit (Fig. 2D). In fixed-cell studies, gH2AX foci (Supplementary Fig.4) appear within 1 hour following drug washout, showing a delayed 53BP1 response in multinucleated cells.

**Fig. 2 -.**
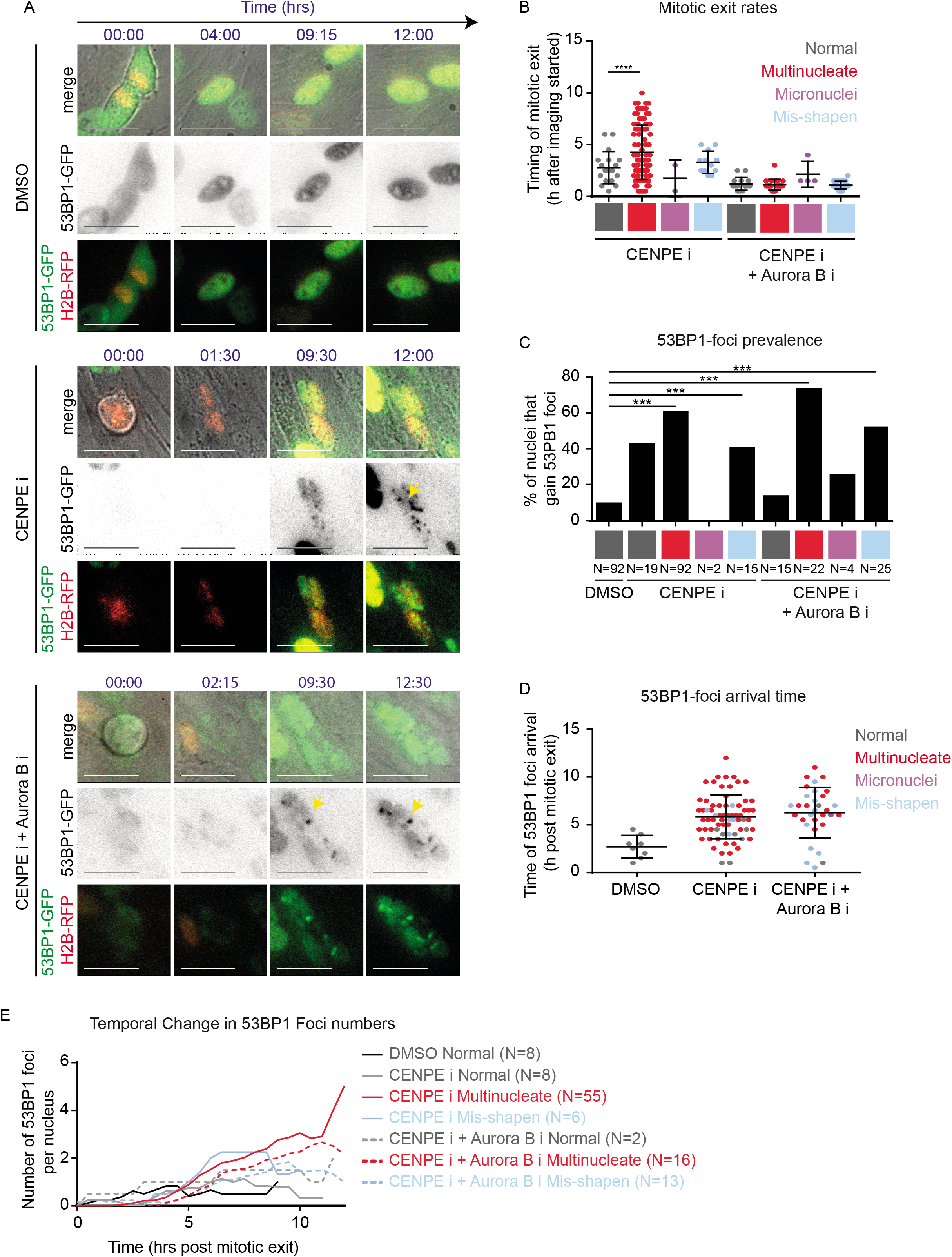
Multinucleate cells exhibit delayed DNA damage signalling and an increase in damage foci through time. A - Representative time-lapse images of normal shaped nuclei exiting mitosis after DMSO treatment, or multinucleated cells exiting mitosis after CENPE i or CENPEi and Aurora Bi treatment as in Sup Fig. 5A. Scale 25μm. Yellow arrows indicate 53BP1-GFP foci within multinucleate cells. In the 53BP1-GFP lane, GFP intensities are inverted to highlight 53BP1-foci as soon as they form (associated supplementary movies present non-inverted GFP intensities). B - Graph shows the timing of mitotic exit, based on nuclear morphology of daughter cells (indicated by colour). Statistical significance was assessed using an unpaired student’s t-test. **** indicates p<0.0001. C - Graph of the proportion of daughter nuclei which gain 53BP1 foci during time-lapse imaging. N values below bars indicate the number of nuclei from at least three independent experimental repeats. Statistical significance was assessed using a proportions test with 95% confidence interval. *** indicates p<0.001. D - Graph of the timing of 53BP1 foci arrival in nuclei which gain 53BP1 foci, after drug treatments as indicated. Time-lapse movies as in A were used to determine the earliest time-point of visible 53BP1 foci following mitotic exit. Each value represents one nucleus. The colour of plotted values represents nuclear morphology. E - Quantification of 53BP1 foci number per nucleus over time from mitotic exit.

In all multinucleated cells, irrespective of slow mitotic slippage or rapid mitotic exit, 53BP1 foci resolution was lacking and the foci number was steadily increasing in late G1 (Supplementary Fig. 5C, Fig. 2E). These single-cell tracking studies reveal that while none of the multinucleated cells resolved 53BP1 foci completely (n=55), 40-50% of normal nuclei displayed 53BP1 foci disappearance indicative of damage clearance (n=8 DMSO and 9 CENPEi cells) (Supplementary Fig. 5C). We conclude that MADD signalling is delayed and MADD remains unresolved.

### MADD-associated 53BP1 remains unresolved despite intact DDR in multinucleated cells

We investigated whether MADD can not be resolved or multinucleated cells are DNA Damage Repair (DDR) deficient by tracking the fate of laser-induced DNA damage in 53BP1-GFP expressing multinucleated cells. Comparing images before and after laser-induced damage showed that cells displaying multinucleation or normal nuclei can rapidly recruit 53BP1-GFP at the laser-damage site (Fig. 3A). The acquisition of laser-induced 53BP1-GFP foci was slightly reduced in number and delayed in multinucleated cells compared to normal cells (Fig. 3B and 3C). Clearance of laser-induced 53BP1 foci was surprisingly slightly more frequent in multinucleated cells compared to normal cells (Fig. 3D). Comparing the rates of laser-induced 53BP1 foci clearance showed no difference between a multinucleated versus normal cell (Fig. 3E and 3F), revealing a proficient DDR pathway in multinucleated cells. Despite DDR proficiency, multinucleated cells did not resolve any of the non-laser induced 53BP1 foci, while normal nuclei resolved 40% of the foci during imaging of up to 15 hours (Fig. 3E). These data show that MADD associated 53BP1 foci cannot be resolved despite a proficient DDR response in multinucleated cells.

**Figure 3 -.**
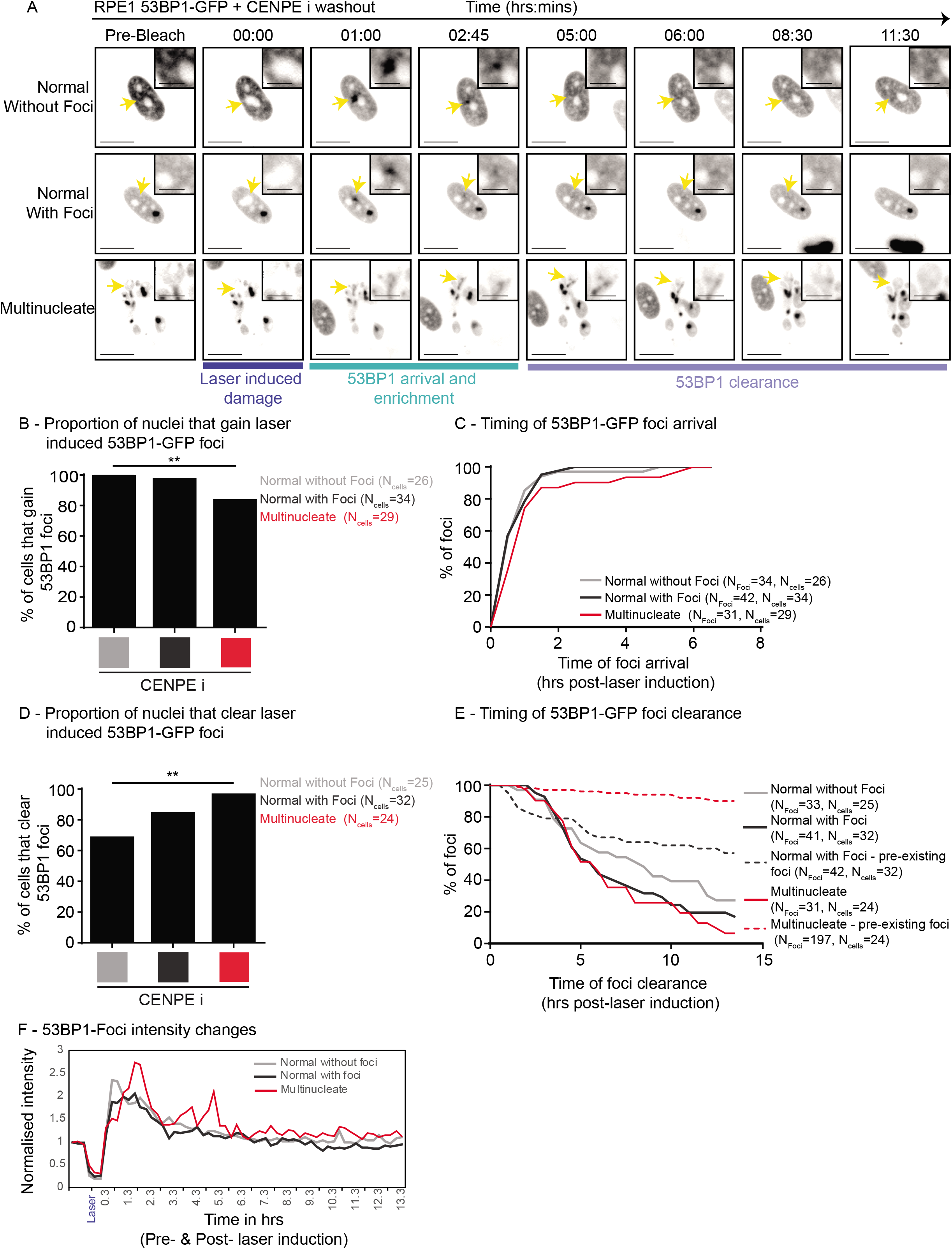
MADD is not resolved despite intact DDR signalling in multinucleate cells. RPE1 H2B-RFP 53BP1-GFP cells were treated with CENPEi for 24 hours, washed and imaging initiated. One laser induced bleach/damage site in each nucleus was tracked. A - Representative pre-bleach, bleach (at 00:00) and postbleach images of nuclei - either normal nuclei without pre-existing 53BP1 foci, normal nuclei with pre-existing 53BP1 foci or multinucleate nuclei with 53BP1 foci. Yellow arrows indicate sites of bleaching and are highlighted in crops. Scale 15μm; insets 5μm. B - Graph shows the proportion of nuclei that gain 53BP1-GFP foci at the site of laser-induced damage, for multinucleate, normal with pre-existing foci and normal nuclei without pre-existing foci. N indicates the number of cells, from across 3 independent repeats. Statistical significance was assessed using a proportions test with a 95% confidence interval. ** indicated p<0.05. C - Graph shows timing, postbleach, of 53BP1-GFP foci arrival at the laser bleach site. D - Quantification of the proportion of nuclei that cleared laser-induced 53BP1-GFP foci, for normal and multinucleate cells. Statistics was assessed using a proportions test with a 95% confidence interval. ** indicates p<0.05. E - Graph shows the timing of 53BP1-GFP foci clearance after bleach time, for foci induced at the bleach site (solid lines) and foci existing prior to laser bleach (dashed lines). N indicates the number of foci and cells analysed. F - Graph shows changes in 53BP1-GFP foci intensity at laser-induced damage site in cells shown in A. 53BP1-GFP intensities were normalised using pre-laser damage intensity values).

### Multinucleation promotes quiescence

Tracking the impact of multinucleation on cell cycle progression, using time-lapse microscopy, revealed an important heterogeneity across multinucleate compartments. In non-transformed hTERT-RPE1 cells expressing mRuby-PCNA (S-phase Marker) and p21-GFP (G1-phase marker), normal nuclei in both CENP-Ei or DMSO treated conditions showed normal appearance and disappearance of PCNA foci (CENP-Ei = 71; DMSO =195 cells; Supplementary Videos 2A, B; Fig. 4A and 4B). In contrast, cells displaying multinucleation showed no PCNA foci formation for at least 14 hours after mitosis, despite the accumulation of nuclear mRuby-PCNA levels in most compartments (114 Cells; Sup Video 2C; Fig. 4B). Consistent with the lack of PCNA foci, multinucleated cells showed an increase in nuclear p21-GFP, 4-5 hours after mitotic exit (Fig. 4B and 4C). Increasing p21-GFP signal was observed in multinucleated cells despite its absence at mitotic exit (Supplementary Video 2C, Fig. 4B), indicating a stress signalling response in late G1. Importantly, a variable increase in the rates and amount of p21 accumulation across different compartments of multinucleated cells was observed (Fig. 4C; n=3 cells). In summary, multinucleation promotes a steady increase in nuclear p21, heterogeneously across compartments, disallowing the onset of DNA replication.

**Figure 4 -.**
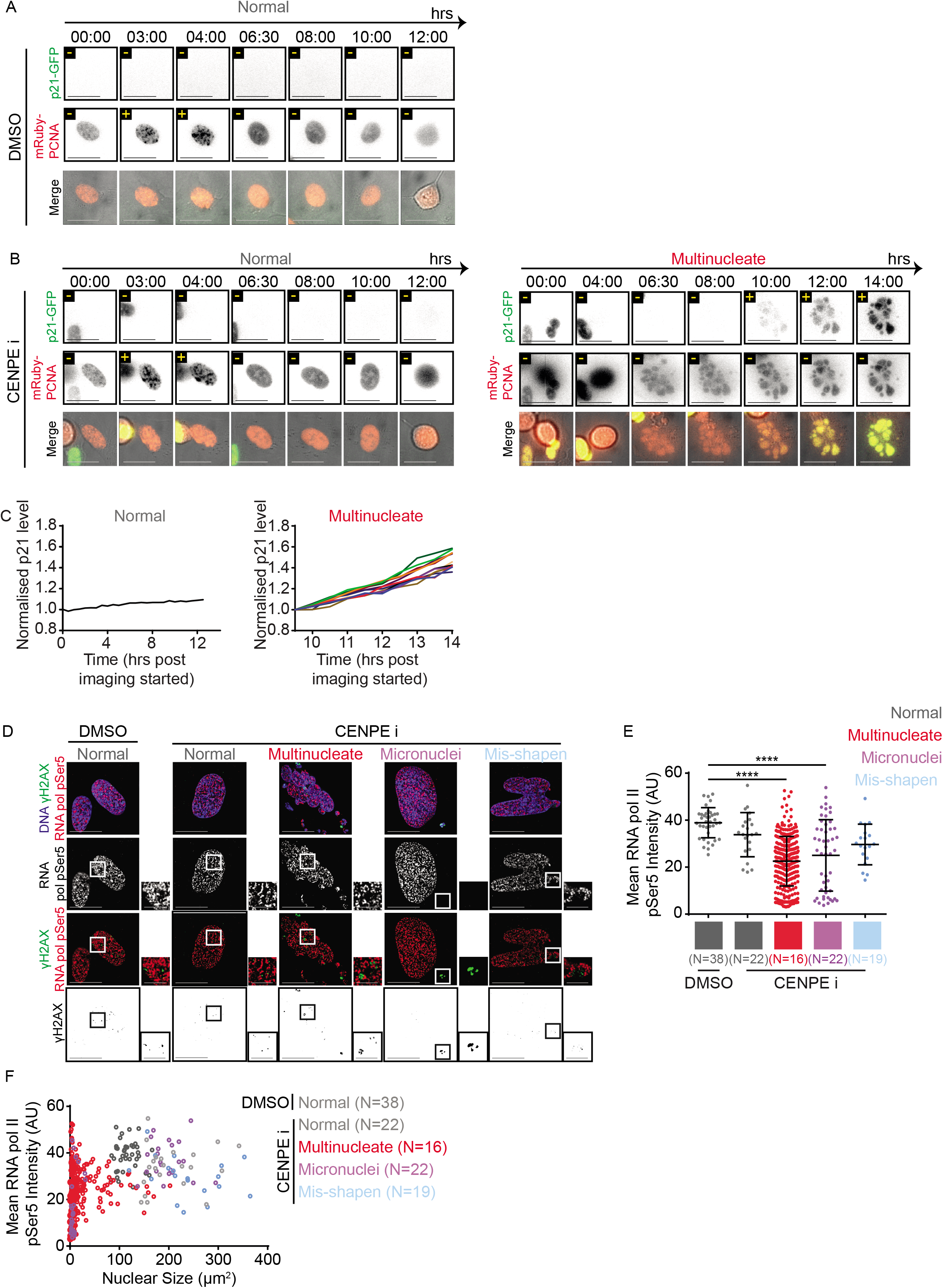
Multinucleation favours heterogeneous nuclear protein levels. A & B - Representative images of RPE1 mRuby-PCNA p21-GFP cells treated with DMSO (A) or CENPEi (B) for 16 hours and washed 10 hours prior to live-cell imaging for 14 hours. Scale bar 25μm. Yellow +/- refers to p21 or PCNA foci positive or negative respectively. Images show a normal shaped nucleus with low p21 and nuclear PCNA foci and a multinucleate cell, exiting mitosis and building p21-GFP levels through time, without gaining nuclear PCNA foci. C - Graph of the mean p21 level per nucleus for the normal, or per compartment for multinucleated cells from movies as in A & B. Values are normalised to the mean p21 value from the first time point measured. PCNA signal was used to identify nuclear areas. Colours of lines correspond to different nuclear compartments. D - RPE1 cells treated with DMSO or CENPEi for 16 hours were fixed 48 hours later for immunostaining with antibodies against gamma H2AX and RNA pol II CTD pSer5. DNA was stained with DAPI. Representative images of non-overlap between gamma H2AX and RNA pol II pSer5 in cells following DMSO or CENPEi treatment. Scale 15μm insets 5μm. E - Graph of the mean RNA pol II CTD pSer5 intensity per nucleus (normal or misshapen nuclei) or nuclear compartment (micronucleated and multinucleate). Each plot represents one nucleus/nuclear compartment respectively. N refers to the number of cells analysed, across 3 independent repeats. Statistical significance was assessed using a one way ANOVA with multiple comparisons. F - Mean RNA pol II CTD pSer5 intensity per nucleus/nuclear compartments plotted against the size of the nucleus/nuclear compartment. Line colours represent nuclear morphology as indicated. N refers to the number of cells from 3 independent repeats.

We hypothesised that if multinucleated cells are not dead and multinucleation disrupts genome-wide function, we should see signs of altered transcription, and we were curious to see if the transcription can occur at MADD-sites. To test whether MADD heterogeneously prevents the function of transcriptional machinery, we immunostained using antibodies against gH2AX and either RNA-Pol II pSer5 (initiation marker) or RNA-Pol II pSer2 (elongation marker). Both markers were present irrespective of nuclear shapes (Fig. 4D; Supplementary Fig. 6C) showing the ability to localise transcriptional machinery and phosphorylate key sites to enable transcriptional initiation and elongation despite large-scale fragmentation of nuclear compartments in multinucleated cells. However, transcription-associated foci did not overlap with gH2AX demonstrating a shutdown of transcription proximal to MADD sites (Fig. 4D; Supplementary Fig. 6C), similar to transcription exclusion at 53BP1 foci arising from replication stress^26^.

Quantifying RNA-Pol II pSer5 or pSer2 foci intensities in nuclei or within a nuclear compartment revealed a statistically significant decrease in mean intensities in some compartments of both multinucleated and micronucleated cells (Fig. 4E; Supplementary Fig. 6D). To investigate whether RNA-Pol intensities are dependent on nuclear compartment size, we correlated changes in mean intensity and compartment size. Micronucleated cells had reduced mean intensities in micronuclei but not primary nuclei, while the compartments in multinucleated cells displayed a unique heterogenous spread of intensities (Fig. 4F; Supplementary Fig. 6E). We tested whether reduced RNA-Pol intensities reflect a reduction in RNA-Pol at each transcriptional site or due to a reduction in the number of transcriptional sites. The proportion of nucleus occupied by RNA-Pol pSer5 or pSer2 foci was reduced in micronucleated and some smaller compartments of multinucleated cells (Supplementary Fig. 6A and 6F). In smaller multinucleate compartments (<40um2) a transcriptional variability is accounted for by a variation in the number of transcriptional sites (Supplementary Fig. 6B and 6G), revealing a nuclear compartment size cut-off for transcription in multi-nucleated cells. These data show exclusion of transcription at MADD sites and a size-dependent reduction in transcription in multinucleated and micronucleated compartments. In summary, the studies of RNA-pol phosphorylation provide three key insights: (i) multinucleated cells are not dead and are transcriptionally active; (ii) multinucleation induces heterogeneity within nuclear compartments with respect to transcriptional initiation and elongation and (iii) transcriptionally active regions are clearly excluded from MADD sites, indicating the wide impact of MADD to the nearby genome.

Because multinucleated cells cannot resolve MADD (Fig. 3) and display increasing p21 (Fig. 4), we tested whether multinucleated cells would remain arrested at G1-S or enter quiescence. To address this, we allowed multinucleated cells to grow for 2 days and assessed DNA damage signalling and cell cycle markers, p53 and phospho-Rb (pRb), a proliferation marker hypophosphorylated in quiescent cells. p53 levels increase in CENPEi-treated cultures as expected following a prolonged mitotic arrest and DNA damage (Supplementary Fig. 1B) and cells were predominantly negative for pRb, unlike controls (Fig. 5A).-To investigate whether entry into quiescence is varied across different forms of atypia, we performed single-cell studies, 2 days after release from mitotic arrest. The proportion of pRb-positive cells was reduced in general but significantly in multinucleated cells (Fig. 5B and 5C). Under conditions inducing DNA damage, a transient reduction in p53 is sufficient to revert quiescent cells into the proliferative state^27,28^. We tested whether reducing p53 will allow multinucleated cells to overcome quiescence and transition into the cell cycle. While there was an increase in total pRB levels following p53 knockdown (Compare CENP-Ei associated lanes of Fig. 5A and 5D), single-cell studies revealed that this increase in pRB is not prominent in multinucleated cells compared to cells displaying other nuclear atypia (Fig. 5E and 5F). We conclude that unlike other nuclear atypia, multinucleation promotes quiescence despite compromised p53 (Fig. 5G).

**Figure 5 -.**
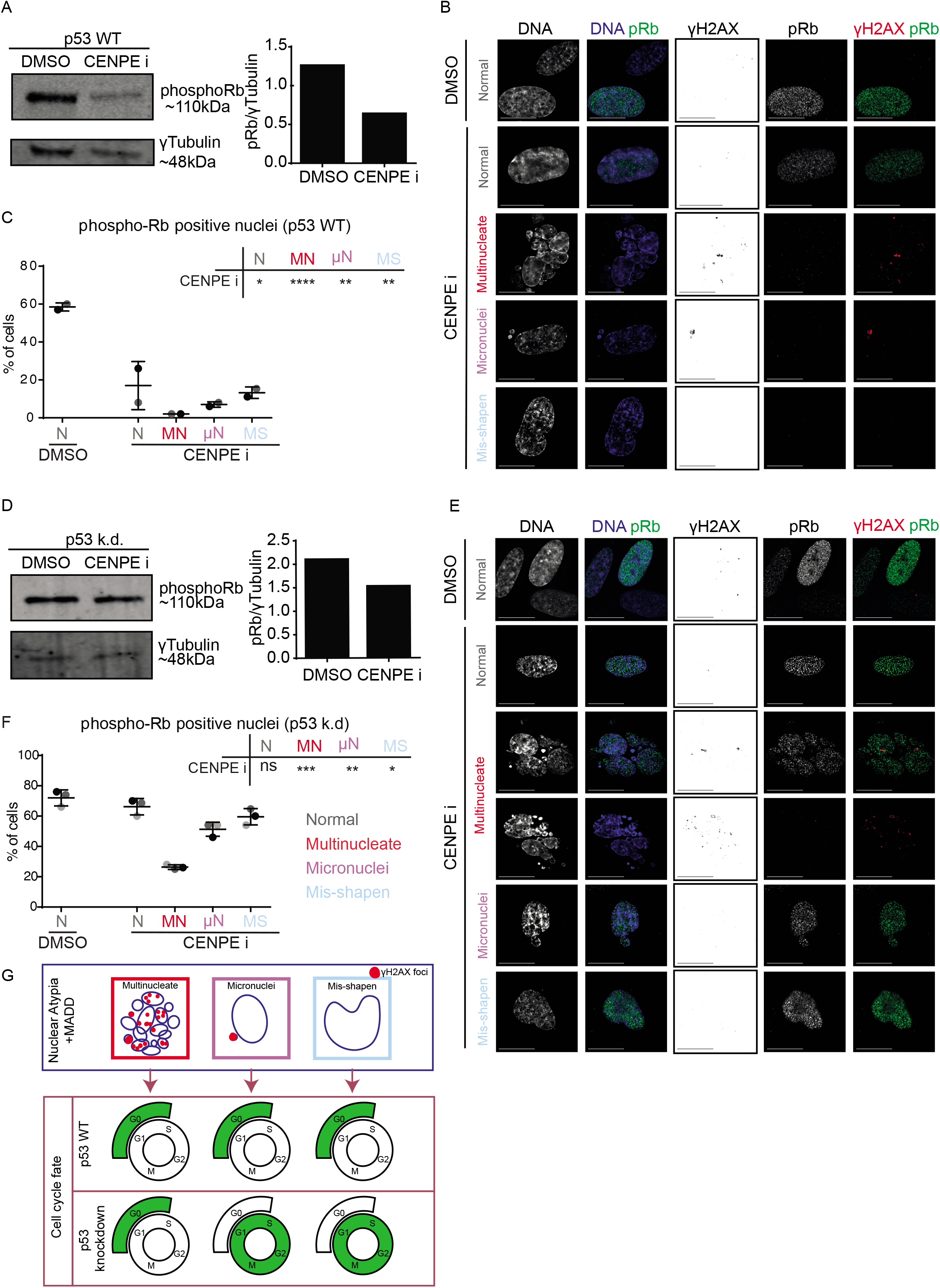
Multinucleation promotes quiescence. RPE1 p53 WT (A-C) or RPE1 H2B-GFP p53 k.d. (D-F) cells were treated with DMSO or CENPEi for 16 hours and 48 hours later cells were immunostained with antibodies against pRb and gamma H2AX, or cells were lysed for immunoblot. A & D - Immunoblot shows pRb or gamma-tubulin levels in RPE1 p53 WT (A) or p53 kd (D) cells following DMSO or CENPEi treatment, as indicated. Note the gamma-tubulin is the same as displayed in Sup Fig 1. Right panel shows a graph of pRb fluorescent intensity, normalised to gamma-tubulin. B & E - Representative images of nuclear atypia following CENPEi treatment of RPE1 p53 WT (B) or p53 kd (E)cells. Scale 15μm. C & F-Graph of proportion of pRb positive or negative RPE1 WT (C) or p53 kd (F) cells, after DMSO or CENPEi treatment. N>100 for WT or >150 for p53 k.d cells from 2 or 3 independent repeats (shown as shades of grey), respectively. Statistical analysis using multiple unpaired t-tests, comparing each morphology after CENPEi treatment to normal nuclei after DMSO. G - Model comparing nuclear atypia shows large-scale DNA damage in multinucleate cells, but not in misshapen nuclei, and in micronucleated cells, gamma H2AX foci are majority confined to the micronucleus. Nuclear atypia causes G0 arrest in p53 WT. In p53 k.d. conditions, only multinucleate cells are G0 arrested.

## Discussion

We report that multinucleated cells are robustly driven to quiescence, despite compromised p53, unlike other forms of nuclear atypia, including micronuclei. This protective aspect of multinucleated cells in blocking DNA replication and S-phase is strongly linked to (i) increasing DNA damage associated 53BP1 bodies in G1-phase which remain unresolved despite an intact NHEJ pathway (ii) heterogeneous nuclear protein localisation and function across compartments (RNA transcription and protein levels), (iii) lack of overlap between DNA damage associated gH2AX and transcriptional initiation or elongation foci and (iv) increasing p21 following mitotic exit, unlike quiescence signalling that normally precedes mitosis in G2^29,30^. We propose this combination of dysfunctional sub-cellular events associated with multinucleation uniquely stabilises p53 function, presenting a protective quiescence phenotype unlike other nuclear atypia that progress through the cell cycle.

Mitotic error induced cell cycle arrest is known, yet a heterogeneity in their cell fates remains unexplained^31–33^; resolving this is important for disease stratification and targeted treatment. To address this gap, we explore heterogeneity in proliferative fate following the same mitotic perturbance. Previously, classes of nuclear atypia have not been quantitatively compared, other than studies of micronuclei alone^3–5^. Prolonged mitosis^34–37^ and chromosome missegregation^20,21,38^ are well-established causes of p53-mediated G1-arrest. Beyond these, we reveal nuclear compartment status as a predictor of proliferative fate following mitotic error. We demonstrate that multinucleation is associated with a heterogeneous change in transcription and protein accumulation across nuclear compartments, and unresolved MADD associated 53BP1 bodies in G1. Surprisingly, MADD sensing, NHEJ signalling, laser-induced DDR and G1-S inhibition operate normally in multinucleated cells. Unresolved 53BP1 bodies associated with MADD may be similar to DSB-SCARS^39^ except that we observe RIF1 accumulation and gH2AX foci and dynamic changes to some (but not all) of the 53BP1 foci in multinucleated cells. The precise molecular reason for differences in 53BP1 foci resolution is unclear and would require future studies downstream of gH2AX-53BP1-RIF1. Nevertheless, the impact of multinucleation on some but not all subcellular processes presents an advantage for targeted treatment of cancers with multinucleated cells.

Micronuclei with cytoplasmic DNA mount a protective immune response^40,41^, but allow proliferation despite DNA damage, in the face of compromised p53. However, multinucleation blocks G1-S despite compromised p53 and presents exposed DNA, offering double-protection. These may be relevant to nuclear atypia induced by cytoskeletal tension (reviewed in^42^), cell-substrate stiffness^43^ or nuclear envelope defects (reviewed in^44^). We propose that unlike other forms of nuclear atypia that show unrestrained proliferation and associated further accumulation of DNA instability, multinucleation presents a protective form which should be considered during pathology assessment and post-treatment with anti-mitotics.

## Supplementary Figures

**Supplementary Figure 1 -.**
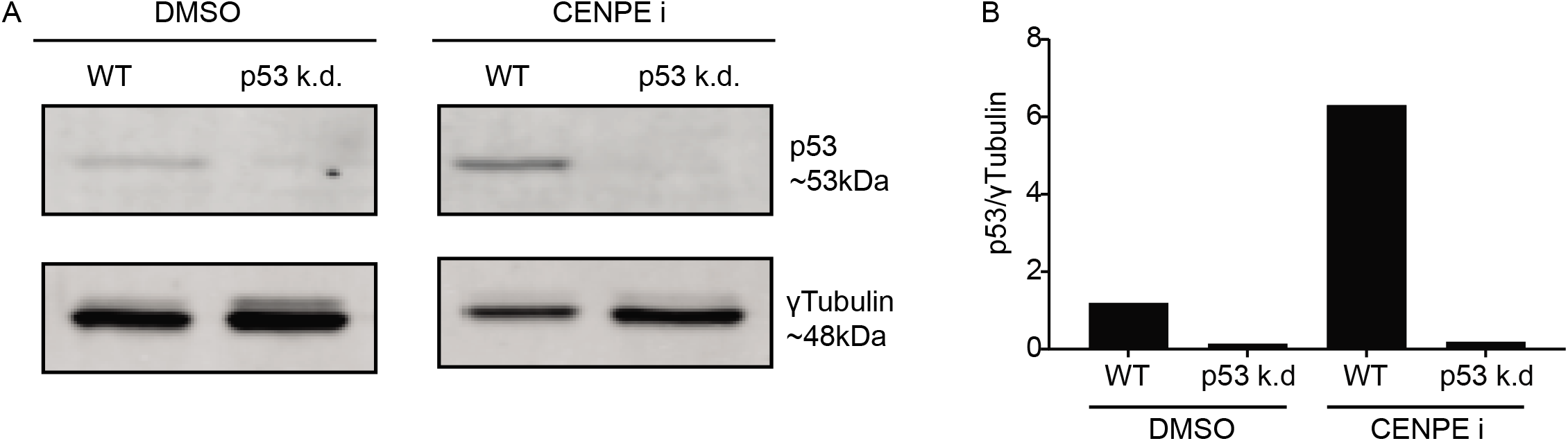
Immunoblot evidence of p53 knockdown. RPE1 p53 WT and RPE1 H2B-GFP p53 k.d. cells were treated with either DMSO (solvent control) or CENPE inhibitor for 16 hours. Drug treatments were then washed out and cells were lysed 2 days later. A - Lysates were run of SDS-PAGE gel and immunoblotted with antibodies against p53 and gamma-tubulin. B - Quantification of immunoblot bands; p53 band intensities were normalised to the intensity of gammatubulin bands.

**Supplementary Figure 2 -.**
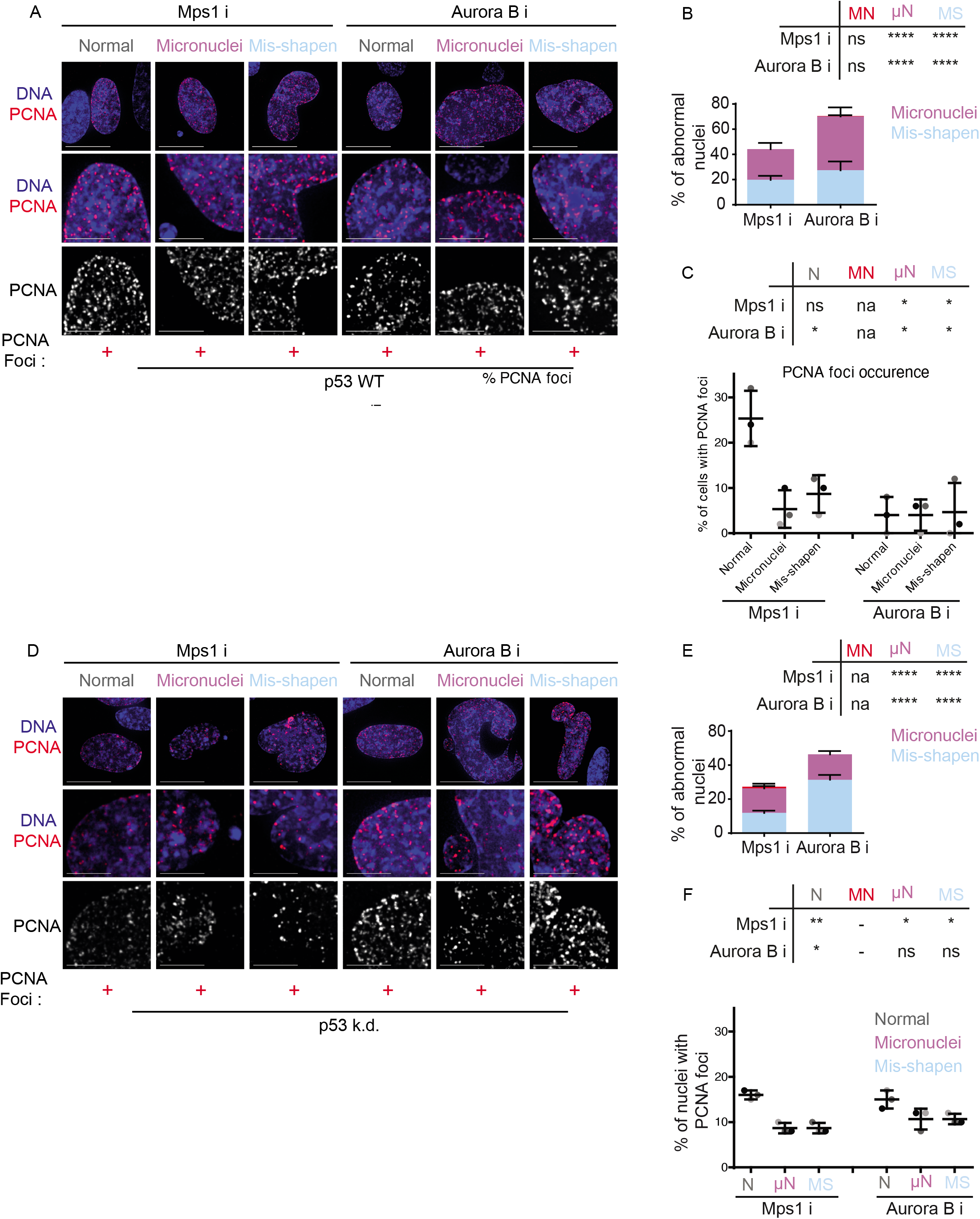
Mis-shapen and micronucleated cells have similar cell cycle responses following an accelerated mitotic progression. RPE1 p53 WT or RPE1 H2B-GFP p53 k.d. cells were treated with MPS1 inhibitor or Aurora B inhibitor for 16 hours then washed out. 48 hours later cells were fixed and immunostained with an antibody against PCNA and DNA was stained with DAPI. A - Representative images of RPE1 p53 WT cells treated with MPS1 or Aurora B inhibitor. Scale bar 15μm, insets 5μm. B - Quantification of nuclear morphology changes following indicated treatments in RPE1 p53 WT cells. N=600 cells from across 3 independent repeats. Statistical analysis was carried out using a two-way ANOVA with multiple comparisons and a confidence interval of 95%. C - Quantification of the percentage of PCNA-foci positive cells, within each nuclear morphology bin, following MPS1 or Aurora B inhibition. N= at least 150 nuclei from across 3 independent repeats shown as shades of grey. Statistical analysis was carried out using multiple unpaired t-tests, comparing each morphology after MPS1 or Aurora B inhibition to normal nuclei after DMSO treatment. D - Representative images of RPE1 H2B-GFP p53 k.d. cells treated with MSP1 or Aurora B inhibitor. Scale bar 15μm, insets 5μm. E - Quantification of nuclear morphology changes following MSP1 or Aurora B inhibitor treatment in RPE1 H2B-GFP p53 k.d. cells. N=900 cells from across 3 independent repeats. Statistical analysis was carried out using a two-way ANOVA with multiple comparisons and a confidence interval of 95%. F - Quantification of the percentage of PCNA-foci positive cells, within each nuclear morphology bin, following MPS1 or Aurora B inhibition. after MPS1 or Aurora B inhibition. N= at least 150 nuclei from across 3 independent repeats shown as shades of grey. Statistical analysis was carried out using multiple unpaired t-tests, comparing each morphology after MPS1 or Aurora B inhibition to normal nuclei after DMSO treatment.

**Supplementary Figure 3 -.**
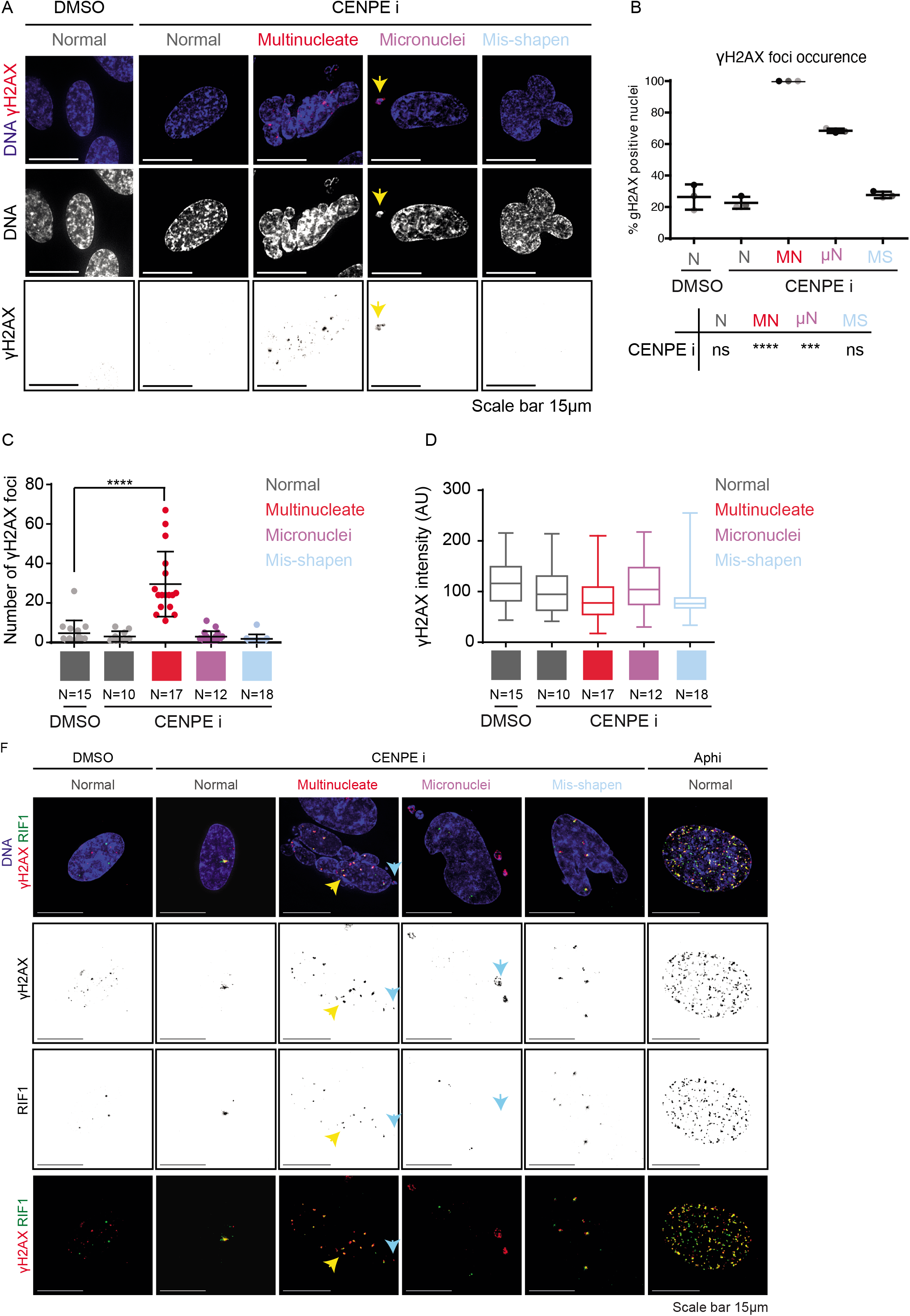
Multinucleation induces DNA damage, increased nuclei with gamma H2AX foci and increased foci within each nucleus. RPE1 cells were treated with DMSO or CENPE inhibitor for 16 hours, then washed out and fixed 2 days later. Cells were immunostained with antibodies against γH2AX and DNA was stained with DAPI. A - Representative images of each nuclear morphology after CENPE inhibition or DMSO treatment. Images are maximum z projections of at least 4 images spanning 2μm. Scale bar 15μm. Yellow arrow indicates a micronucleus. B - Graph to show the percentage of each nuclear morphology which had γH2AX foci within the nucleus, following DMSO or CENPE inhibition. N=300 nuclei per morphology from across 3 independent repeats shown as shades of grey. Significance was assessed with multiple unpaired t-tests to compare each nuclear morphology after CENPE inhibition to normal nuclei after DMSO treatment. C - Quantification of the number of γH2AX foci per nucleus, in nuclei with γH2AX foci. Quantification is divided based upon nuclear morphology after DMSO or CENPE inhibition. N indicated the number of cells, from across 3 independent repeats. Significance was assessed using an unpaired t-test. Other nuclear morphologies were non-significant when compared to normal nuclei after DMSO treatment. D - Quantification of the γH2AX fluorescence intensity per foci, in nuclear morphologies indicated after DMSO or CENPE inhibitor treatments. Boxes represent second and third quartiles respectively, middle line displays median, and whiskers span minimum to maximum values. N indicated the number of cells analysed, taken across 3 independent repeats. F - RIF1 localises to sites of DNA damage in multinucleate cells, but not inside micronuclei. RPE1 cells were treated with DMSO or CENPE inhibitor for 16 hours then washed out and 2 days later fixed and immunostained with antibodies against gamma H2AX and RIF1. DNA was stained with DAPI. As a positive control RPE1 cells were treated with Aphidicolin for the duration of the experiment. Representative images of cells with differing nuclear morphology after indicated drug treatments; Normal after DMSO (N=29 cells), Normal (N=8), Multinucleate (N=12), Micronuclei (N=8) and Mis-shapen nuclei (N=7) after CENPE inhibition, and normal nuclei after Aphidicolin treatment (N=8). Blue arrow indicates gamma H2AX foci but no RIF1 localisation within micronucleus, and yellow arrow indicates site of gamma H2AX and RIF1 co-localisation. Scale bar 15μm.

**Supplementary Figure 4 -.**
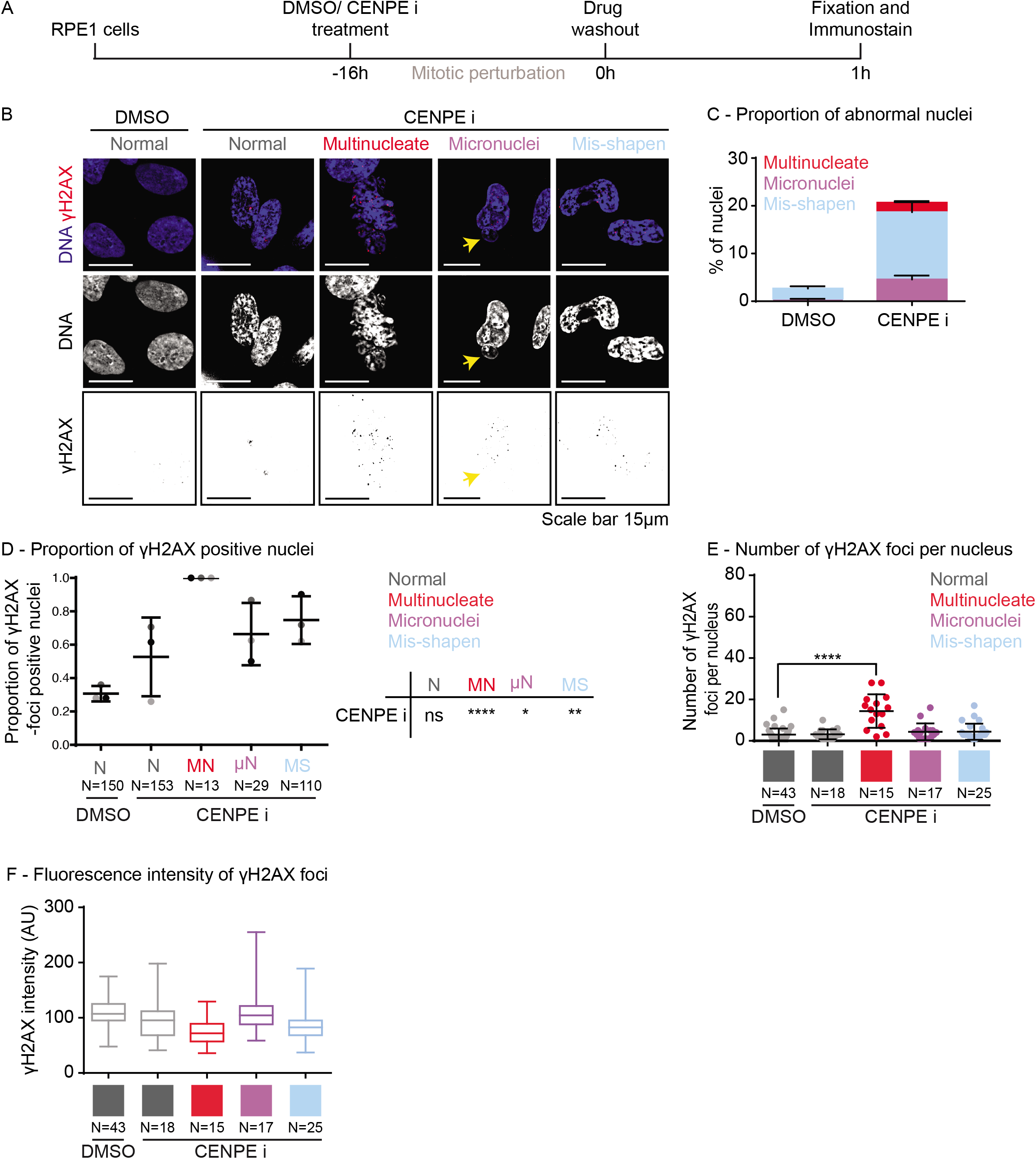
Multinucleate cells have DNA damage 1-hour postdrug washout. A - Experimental regime; RPE1 cells were treated with CENPE inhibitor or DMSO for 16 hours, the drug then washed out and cells were fixed 1 hour later. Cells were immunostained with antibodies against γH2AX and DNA was stained with DAPI. B - Representative images of nuclei as treated as indicated in A. Scale bar 15μm. Yellow arrow indicates a micronucleus. C - Quantification of the proportion of cells with abnormal nuclei after treatment as described in A. N=900 cells per condition, taken from 3 independent repeats. D - Quantification of the proportion of each nuclear abnormality with γH2AX foci within the nucleus. N is indicated below bars, taken from 3 independent repeats. Statistical analysis was carried out using multiple unpaired t-tests, comparing each morphology after CENPE inhibition to normal nuclei after DMSO treatment. F - Quantification of the number of γH2AX foci per nucleus, for each nuclear morphology, after DMSO or CENPE inhibitor treatment. N is indicated under each bar, taken from 3 independent repeats. Significance was assessed using an unpaired t-test. Other nuclear morphologies were non-significant when compared to normal nuclei after DMSO treatment. G - Quantification of the fluorescence intensity of γH2AX, per foci, in nuclear morphologies indicated after DMSO or CENPE inhibitor treatment. Boxes represent second and third quartiles respectively, middle line displays median and whiskers span minimum to maximum values. N indicated the number of cells analysed, taken across 3 independent repeats.

**Supplementary Figure 5 -.**
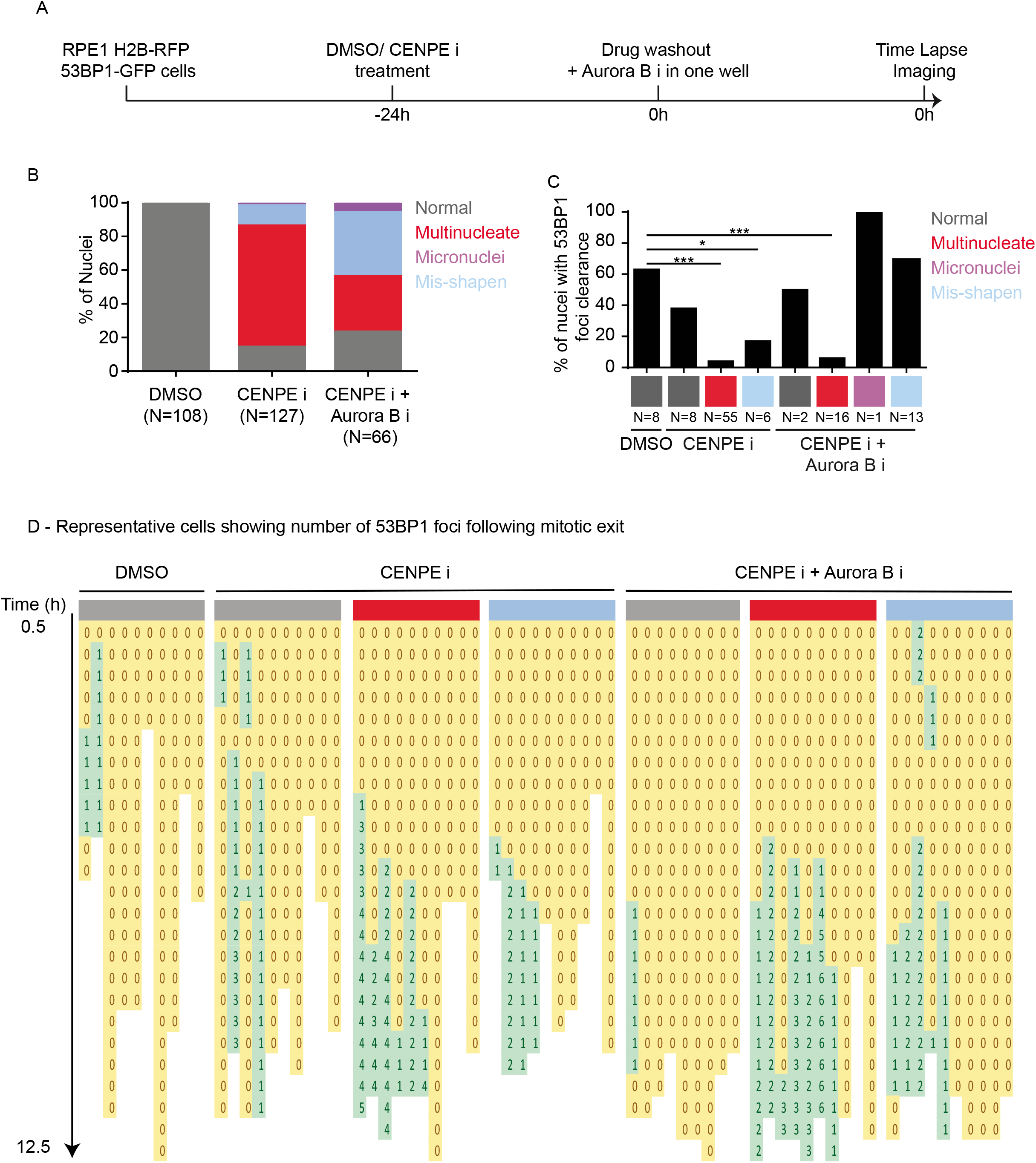
Individual tracking of 53BP1-GFP foci dynamics. A - Experimental regime; RPE1 H2B-RFP 53BP1-GFP cells were treated with DMSO or CENPE inhibitor for 24 hours, then washed out and one well additionally treated with Aurora B inhibitor. Cells were then imaged for at least 12 hours with an image every 30 minutes. B - Quantification of the nuclear morphology of daughter cells exiting mitosis after DMSO, CENPE inhibitor or CENPE inhibitor plus Aurora B inhibitor treatment. N, as indicated, is the number of daughter nuclei. C - Quantification of the proportion of nuclei which display clearance of 53BP1 foci during time-lapse imaging. N values below bars indicate the number of nuclei. Statistical significance was assessed using a proportions test, with a 95% confidence interval. *** indicates p<0.001, * indicates p<0.05. D - Representative tracks of individual cells with the number of 53BP1 foci over time following mitotic exit. The number and colour represent the number of 53BP1 foci per nucleus.

**Supplementary Figure 6 -.**
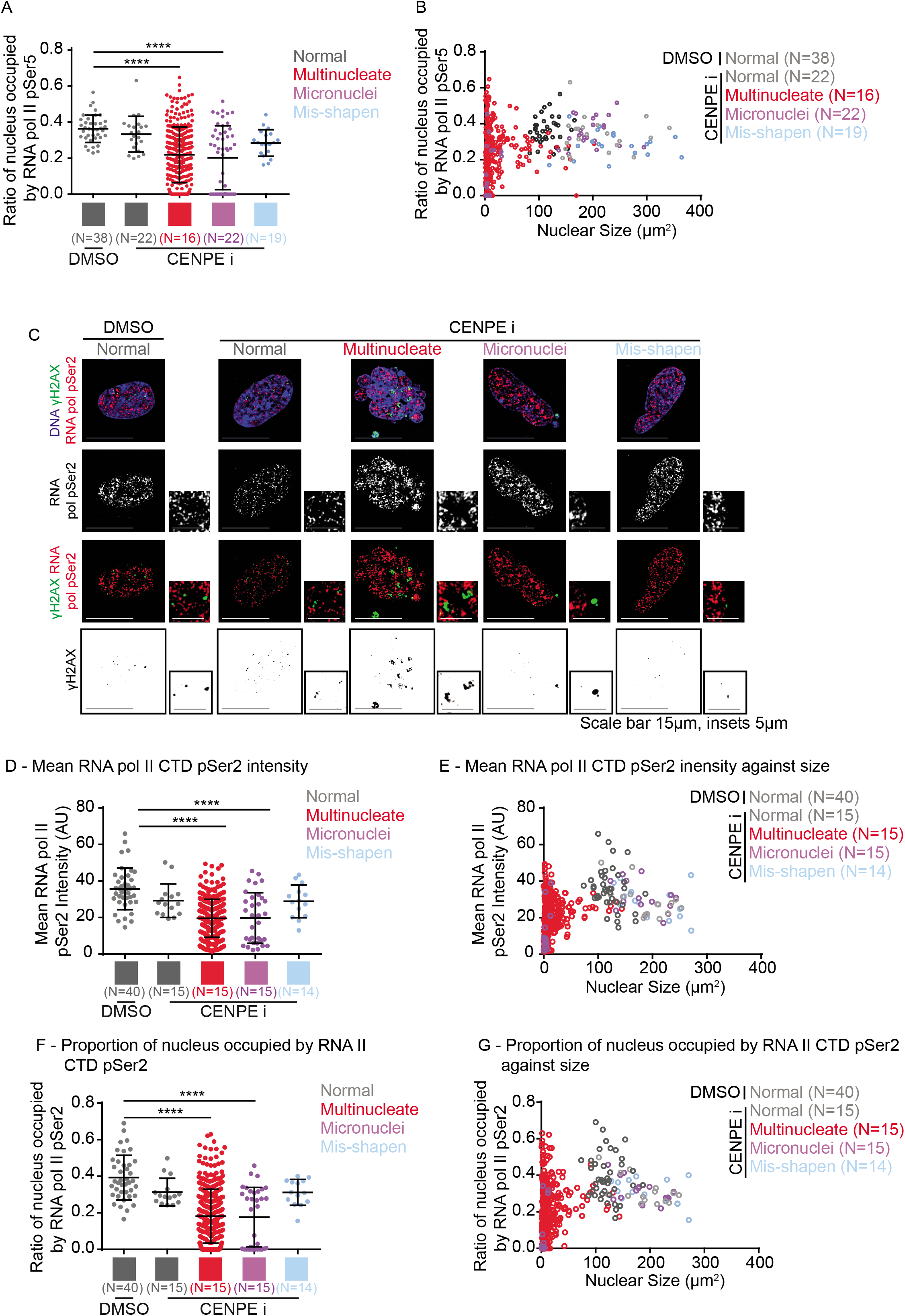
Transcription initiation and elongation variable in smaller compartments of multinucleate cells. A - Quantification of the proportion of the nucleus occupied by RNA pol II CTD pSer5, for each nucleus (normal and misshapen nuclei) or nuclear compartment (micronuclei and multinucleate). N values indicate the number of cells analysed. Statistical significance was assessed using a one way ANOVA with multiple comparisons. B - Plot of the proportion of the nucleus occupied by RNA pol II CTD pSer5 against nucleus or nuclear compartment size. Colours of plots represent nuclear morphology as indicated. N values indicate the number of cells. C-G - RPE1 cells were treated with DMSO or CENPE inhibitor for 16 hours, then washed out and fixed 2 days later. Cells were immunostained with antibodies against γH2AX and RNA polymerase II CTD pSer2 and DNA was stained with DAPI. C - Representative images of differing nuclear morphologies after CENPE inhibitor treatment, and normal nuclei after DMSO treatment. Scale bar 15μm, insets 5μm. D - Quantification of the average RNA pol II CTD pSer2 intensity per nucleus (normal or misshapen nuclei) or nuclear compartment (micronucleated and multinucleate). Each plot represents one nucleus/nuclear compartment respectively. N indicated below the bar is the number of cells analysed, from across 3 independent repeats. Statistical significance was assessed using a one way ANOVA with multiple comparisons. E - Average RNA pol II CTD pSer2 intensity per nucleus/nuclear compartments plotted against the size of the nucleus/nuclear compartment. Colours of plots represent nuclear morphology as indicated. N, as indicated, is number of cells taken from across 3 independent repeats. F - Quantification of the proportion of the nucleus occupied by RNA pol II CTD pSer2, for each nucleus (normal and misshapen nuclei) or nuclear compartment (micronuclei and multinucleate). N values indicate the number of cells analysed. Statistical significance was assessed using a one way ANOVA with multiple comparisons. G - Graph of the proportion of the nucleus occupied by RNA pol II CTD pSer5 against nucleus or nuclear compartment size. Colours of plots represent nuclear morphology as indicated.

## Supplementary Videos

**Supplementary Videos 1 - Multinucleate cells gain 53BP1-GFP foci in G1.** RPE1 H2B-RFP 53BP1-GFP cells were treated with DMSO or CENPE for 24 hours. Cells were then washed and one well was additionally treated with Aurora B inhibitor. Cells were then imaged for at least 12 hours using time-lapse microscopy. Imaging shows 53BP1-GFP (top left), H2B-RFP (top right), widefield (lower left) and merge (lower right) channels. A - Normal daughter nuclei exiting mitosis following DMSO treatment, do not gain 53BP1-GFP foci during the course of the imaging. B - A CENPE inhibitor arrested cell exits mitosis into a multinucleate daughter cell which gains initial 53BP1-GFP foci at 08:30hrs and increases in number until the end of the Video. Corresponding images in Figure 5 (first two panels) present intensity adjustments to rule out any low intensity foci in early G1. C - CENPE inhibitor arrested mitotic cell is released from mitosis with Aurora B inhibition and forms a multinucleate daughter cell. Numerous 53BP1-GFP foci appear at 09:00hrs which increase in number until the end of imaging.

**Supplementary Videos 2 - Nuclear PCNA foci in normal nuclei after DMSO or CENPE inhibition but not in multinucleate cells, which have increasing p21 levels.** RPE1 mRuby-PCNA p21-GFP cells were treated with DMSO (A) or CENPE inhibitor (B, C) for 16 hours. Drugs were then washed out and live-cell imaging started 10 hours later. Imaging shows p21-GFP (top left), mRuby-PCNA (top right), widefield (lower left) and merge (lower right) channels. A - Normal nucleus after DMSO treatment shows nuclear PCNA foci and lack of p21. B - Normal nucleus after CENPE inhibition shows nuclear PCNA foci and lack of p21. C - Multinucleate cell after CENPE inhibition exits mitosis and builds p21 level through time, without gaining nuclear PCNA foci. Scale bar 25μm. Videos correspond to images shown in Figure 2.

## Materials and Methods

### Cell culture and cell lines

RPE1 cells were cultured in DMEM F12 supplemented with 10% fetal calf serum and penicillin and streptomycin. For experiments they were plated onto 13mm round coverslips for immunofluorescence or onto glass-bottomed dishes (LabTech) for live-cell imaging. All live-cell studies were conducted in stable cell lines: RPE1 H2B-RFP 53BP1-GFP, RPE1 H2B-GFP p53 k.d, mRuby-PCNA and p21-GFP cell lines were all grown in DMEM-F12 and filmed in L15 media.

### Drug treatments

To disrupt mitosis cells were treated with 10nM GSK923295 to inhibit CENPE, 1uM Reversine to inhibit Mps1 and 10uM ZM447439 to inhibit Aurora B. For immunofluorescence experiments drug treatments were incubated for 16 hours then washed off and fixed 48 hours later, unless indicated otherwise in text.

### Live-cell time-lapse imaging

To image cell cycle progression in RPE1 p21-GFP mRuby-PCNA cells, cells were treated with 10nM GSK923295 or DMSO control for 16 hours. Cells were then washed and then additionally treated with DMSO or 10uM ZM447439. 10 hours later cells were washed and media changed to Leibovitz (L15) imaging media at 37°C and imaging started. An image was taken at least every 30 minutes for 14 hours. Images were acquired using a 40x 0.75NA objective on a DeltaVision Core^TM^ microscope (GE Healthcare) with a Cascade2 camera under EM mode.

To image 53BP1-GFP foci arrival in cells following mitotic disruption RPE1 H2B-RFP 53BP1-GFP cells were treated with DMSO or GSK923295 for 24 hours. Cells were washed and then transferred to 37°C Leibovitz imaging media with DMSO or ZM447429 in the media. Cells were then imaged for at least 12 hours with an image taken at least every 30 minutes. Images were acquired using a 40x 0.75NA objective on a DeltaVision Core™ microscope (GE Healthcare) with a Cascade2 camera under EM mode.

### FRAP photobleaching to induce 53BP1-GFP foci

The Deltavision Core™ microscope was used with the FRAP tool using Quantifiable laser module components (488nm laser). Target points were identified and then bleached with a pulse duration of 1 second and a laser power of 30%. 3 pre-bleach and 3 post-bleach images were taken at each target site, all 0.5 seconds apart, using a 60x 1.42 NA objective, using CoolSnap HQ camera (Photometrics). Following each site being bleached, time-lapse microscope was initiated for the following 13 hours with an image acquired every 30 minutes, using a 40x 0.75 NA objective with a Cascade2 camera under EM mode.

### Immunofluorescence

For immunofluorescence the following antibodies were used; PCNA (CST, 3586S, 1:1000), Phospho-Rb (Ser 807/811) (CST, 8516S, 1:1000), γH2AX (Abcam, ab26350, 1:800),γH2AX (CST. 1:1000), GFP (Abcam, ab290, 1:1000), RNA polymerase II CTD repeat phospho-Ser2 (Abcam, ab126353, 1:1000) and RNA polymerase II CTD repeat phospho-Ser5 (Abcam, ab5408, 1:1000). DNA was stained with DAPI. Cells were fixed with ice-cold methanol and blocked with 1% BSA in PBS before immunostaining. Images of immunostained cells were acquired using a 100x 1.2 NA objective on a DeltaVision Core microscope with a CoolSnap HQ camera (Photometrics). Cells were scored as positive for PCNA foci when foci signal intensities associated with the PCNA foci (inside the nuclei) were well above the background speckles outside the nuclei.

### Nuclear atypia scoring

DAPI stained nuclei were categorised based upon nuclear appearance, for nuclear morphology. As in Figure 1B, normal nuclei were those that had a continuous nuclear periphery and a regular oval shape, abnormal shaped nuclei were those that deviated from this regular oval shape but the nuclear material was contained within one nuclear compartment. Micronucleated cells have one or a few nuclear compartments external to the primary nucleus, containing one or a fragment of a chromosome. Multinucleate cells have multiple nuclear compartments varying in size, and can include micronuclei.

### Image analysis

Image J was used for measurements of fluorescent intensities, using 8-bit images. Softworx was used for manual image analysis of 53BP1-GFP foci tracking.

### Statistical analysis

Graphs were plotted using Graphpad Prism, which was also used for statistical testing, except form proportions tests which were carried out in Microsoft Excel. In all graphs presented error bars represent standard deviation. In statistical tests presented the following indications for p values were used; non-significant - ns for p>0.05, * for p <0.05, ** for p<0.01, *** for p><0.001, **** for p><0.0001.

## Acknowledgements

The work was supported by funding to Draviam by a BBSRC research grant (R01003X/1) and QMUL start-up grant (SBC8DRA2). We thank the Medema and Barr labs for providing RPE1 H2B-RFP 53BP1-GFP cell line and RPE1 H2B-GFP p53 k.d. cell line as gifts.

## Notes

### Competing Interest Statement

The authors have declared no competing interest.

